# High-resolution positron emission microscopy of patient-derived tumor organoids

**DOI:** 10.1101/2020.07.28.220343

**Authors:** Syamantak Khan, June Ho Shin, Valentina Ferri, Ning Cheng, Julia E. Noel, Calvin Kuo, John B. Sunwoo, Guillem Pratx

## Abstract

Organoid tumor models have found application in a growing array of cancer studies due to their ability to closely recapitulate the structural and functional characteristics of solid tumors. However, organoids are too small to be compatible with common radiological tools used in oncology clinics. Here, we present a microscopy method to image ^18^F-fluorodeoxyglucose in patient-derived tumor organoids with spatial resolution up to 100-fold better than that of clinical positron emission tomography (PET). When combined with brightfield imaging, this metabolic imaging approach functionally mirrors clinical PET/CT scans and provides a quantitative readout of cell glycolysis. In particular, the specific avidity of a tumor for FDG, or lack thereof, was maintained when the tumor cells were grown *ex vivo* as tumor organoids. In addition, cisplatin treatment caused a dose-dependent decrease in the metabolic activity of these organoids, with the exception of one patient whose tumor was also resistant to cisplatin treatment. Thus, FDG-imaging of organoids could be used to predict the response of individual patients to different treatments and provide a more personalized approach to cancer care.

---

Patient-derived tumor organoids are miniature, three-dimensional, self-organized tissue culture models that are derived from primary patient tumor cells and studied in the laboratory^1,2^. These organoid cultures closely recapitulate the genetic and morphological heterogeneity of solid tumors and stromal components present in the original tumor^1-10^. They also retain the immune microenvironment of solid tumors, including the T cell receptor repertoire and PD-1/PD-L1-dependent immune checkpoint status^5^. Patient-derived organoids contain the same cancer mutations and genetic variations that are present in the patient of origin, and thus they can assist in the selection of individualized treatment, especially for patients who fail to respond to first-line therapy ^3,11,12^. Additionally, panels of organoids derived from patient cohorts capture inter-patient variability of tumor phenotype for drug screens^8,13^. They can also be very useful to model rare cancer phenotypes^14^. Finally, organoids have fast turnaround time, high level of control and reproducibility, low cost, and higher throughput, and thus offer an alternative to patient-derived xenografts for certain applications.

While many studies have employed organoids to assess the effects of different therapies, relatively little work has been done in the area of diagnostic imaging. One significant limitation, of course, is that the resolving power of common radiological tools is not suited to the miniature size of organoids. The ability to image organoids using clinical contrast agents with high spatial resolution would facilitate the rapid translation of biomedical information, since the same non-invasive biomarkers could be used in pre-clinical organoid studies and clinical trials. Organoids could also be used as a platform to develop novel clinical contrast agents. This very concept, 15 years ago, drove the development of miniaturized scanners dedicated to small animal imaging, which are now in widespread use at academic research centers and in the labs of pharmaceutical companies^15,16^. However, few non-invasive imaging biomarkers are applicable to both organoids and patients, due to the 1000-fold difference in scale. Several biological endpoints are used for organoid studies, but these bear little relevance to clinical biomarkers and don’t easily translate *in vivo*. For instance, organoid response to treatment can be assessed using colony formation assays^17^, viability assays^18^, and optical redox imaging^19,20^, all of which are far removed from the standard clinical workflow. Likewise, radiological criteria such as RECIST (Response Evaluation Criteria in Solid Tumors)^21^ are challenging to apply to organoid cultures.

The gold standard in cancer clinics for molecular assessment of normal and diseased tissues is positron emission tomography (PET), which is often combined with computed tomography (CT) to achieve multimodal noninvasive imaging of disease *in vivo*. The discovery of the Warburg effect has led to the extensive use of ^18^F-fluorodeoxyglucose (FDG) as the main PET tracer to map glucose metabolism in primary and metastatic tumors^22^. In the last decade, FDG-PET has been shown to drastically improve the diagnosis, staging and subsequent treatment of cancers, and it has emerged as a highly reliable tool for accurate assessment of treatment response after chemotherapy and radiotherapy. Yet, few studies—if any—have investigated the potential of FDG as a biomarker for assessing the response of patient-derived tumor organoids to therapy.

While the low resolution of conventional PET (> 1 mm) is a significant limitation for organoid imaging, recent work has shown the feasibility of imaging PET tracers in 2D cell cultures with single-cell resolution using an approach known as radioluminescence microscopy (RLM). RLM uses a scintillator crystal to convert beta radiation emanating from the cells into optical flashes detectable in a single-photon sensitive microscope, and thus provide high-resolution 2D imaging of radionuclide distribution in live cells^23,24^.

This study presents a proof of principle for imaging organoids using the RLM technique. We call this specific method of imaging as Positron Emission Microscopy of Organoids (oPEM) to highlight its clinical relevance to PET. In combination with brightfield (BF) and fluorescence microscopy, this technique allows multimodal imaging of organoids at higher resolution using fluorescent probes and clinically relevant PET radiotracers. To investigate this approach, patient-derived organoids were grown for 2-3 weeks in 3D hydrogel from tumor tissues derived from head-and-neck and thyroid cancer patients (Supplementary Table 1). Following this, the organoids were treated with ^18^F-FDG and mounted on transparent CdWO_4_ scintillator plates for imaging with RLM (Fig. 1). The spatial distribution of FDG within organoids was superimposed onto standard brightfield and fluorescent micrographs to identify and characterize areas of high glycolysis. The level of FDG uptake of individual organoids was then quantitatively compared to the uptake of the cells in the tumor of origin, as measured *in vivo* using FDG-PET. This analysis revealed a considerable agreement between organoid uptake *in vitro* and tumor uptake *in vivo*. Finally, as a further demonstration of the utility of oPEM, organoids were grown from both cisplatin-sensitive and cisplatin-resistant tumors, and their response to cisplatin was assessed according to the relative difference in FDG uptake.

**Figure 1.**
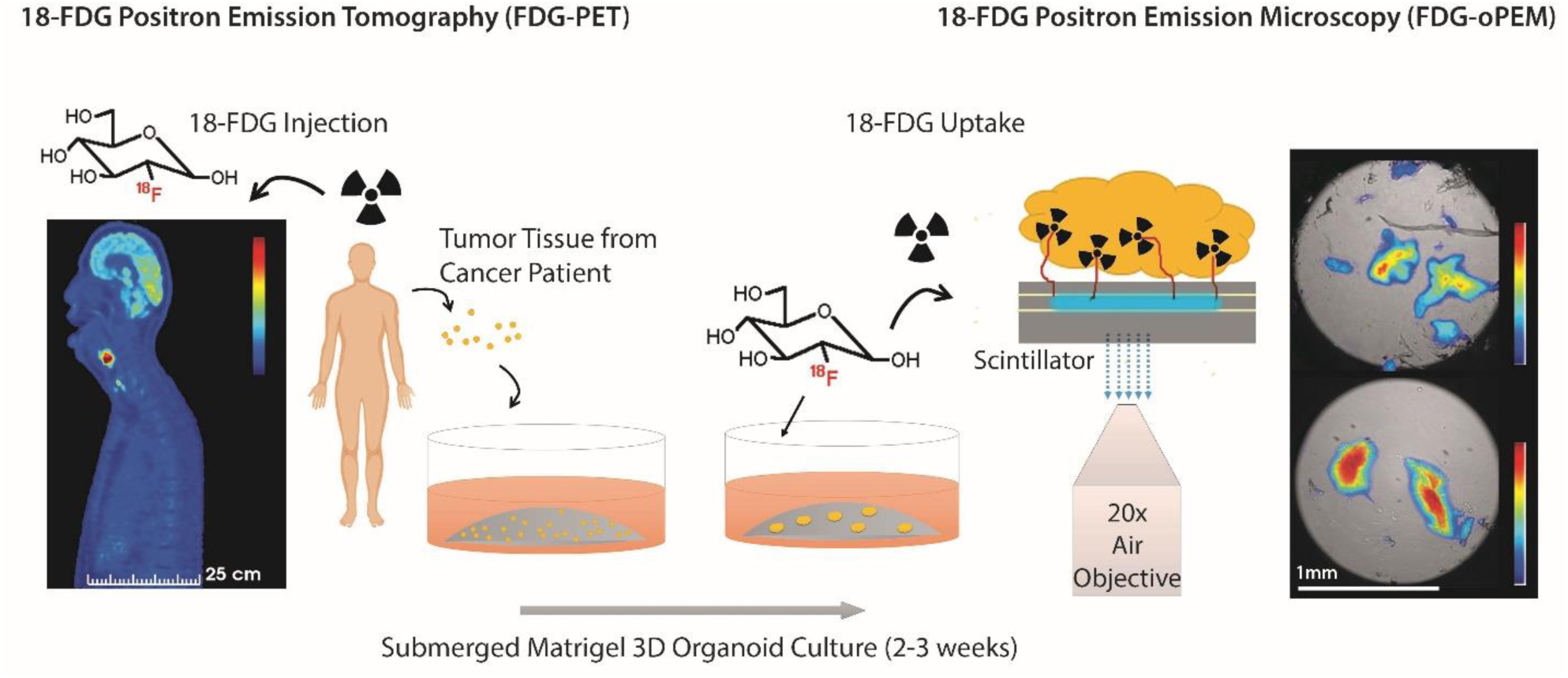
Schematic representation of the workflow. Tumor organoids were cultured for 2-3 weeks in a submerged Matrigel culture system. Patient-derived organoids (right) were imaged with high resolution using FDG, a radiotracer commonly used in the clinic for diagnosis and staging of head-and-neck cancer patients (left). The organoids were mounted on thin inorganic scintillators and the resulting scintillation light was imaged in a highly sensitive microscope through a 20X objective lens. The scale bars of PET and oPEM images highlight the large difference in image resolution.

## Results

### Tumor organoids recapitulate the microenvironment of the original tumor

Tumor organoids were seeded from processed surgical samples of head and neck cancer patients and cultured in basement membrane extract (BME), a soluble form of basement membrane purified from Engelbreth-Holm-Swarm (EHS) tumor. A specialized culture medium (EN medium) containing DMEM/F-12 supplemented with 10% Noggin-conditioned media, Nicotinamide (10 mM), N-Acetylcysteine (1 mM), B-27 minus vitamin A (1X), Pen-Strep (1X), and EGF (50 ng/mL) was used to grow organoids.

These tumor organoids closely recapitulated the most salient features of the tumor of origin. After 3 weeks of culture, they displayed dysplastic epithelial features with keratin production and disorganized growth patterns, typical of squamous cell carcinoma (Fig. 2b & Supplementary Fig. 1). Fluorescent labeling of the nucleus and cell membrane highlighted morphological details of the tumor and stromal components (Fig. 2c), including cancerous tissue (inset, red) and fibroblast-rich stroma (inset, green) around the organoid culture. Overgrowth of the fibroblast population was observed after a few weeks of culture. Fluorescence immunostaining of the organoid and the corresponding tumor of origin highlighted functional and cellular similarities between the two tissues (Fig. 2d, e). In both samples, E-Cadherin selectively identified the cancerous epithelial cells, Vimentin the tumor-associated fibroblast, and CD3 the tumor-infiltrating T-cells. The retention of stromal components after 7 days of *in vitro* culture is a crucial advantage of this cancer model, as treatment response is often influenced by a biologically active tumor stroma and its complex interactions with the cancer cells. In summary, the tumor organoids used in this study are effective and clinically relevant *in vitro* models for studying head-and-neck cancers.

**Figure 2.**
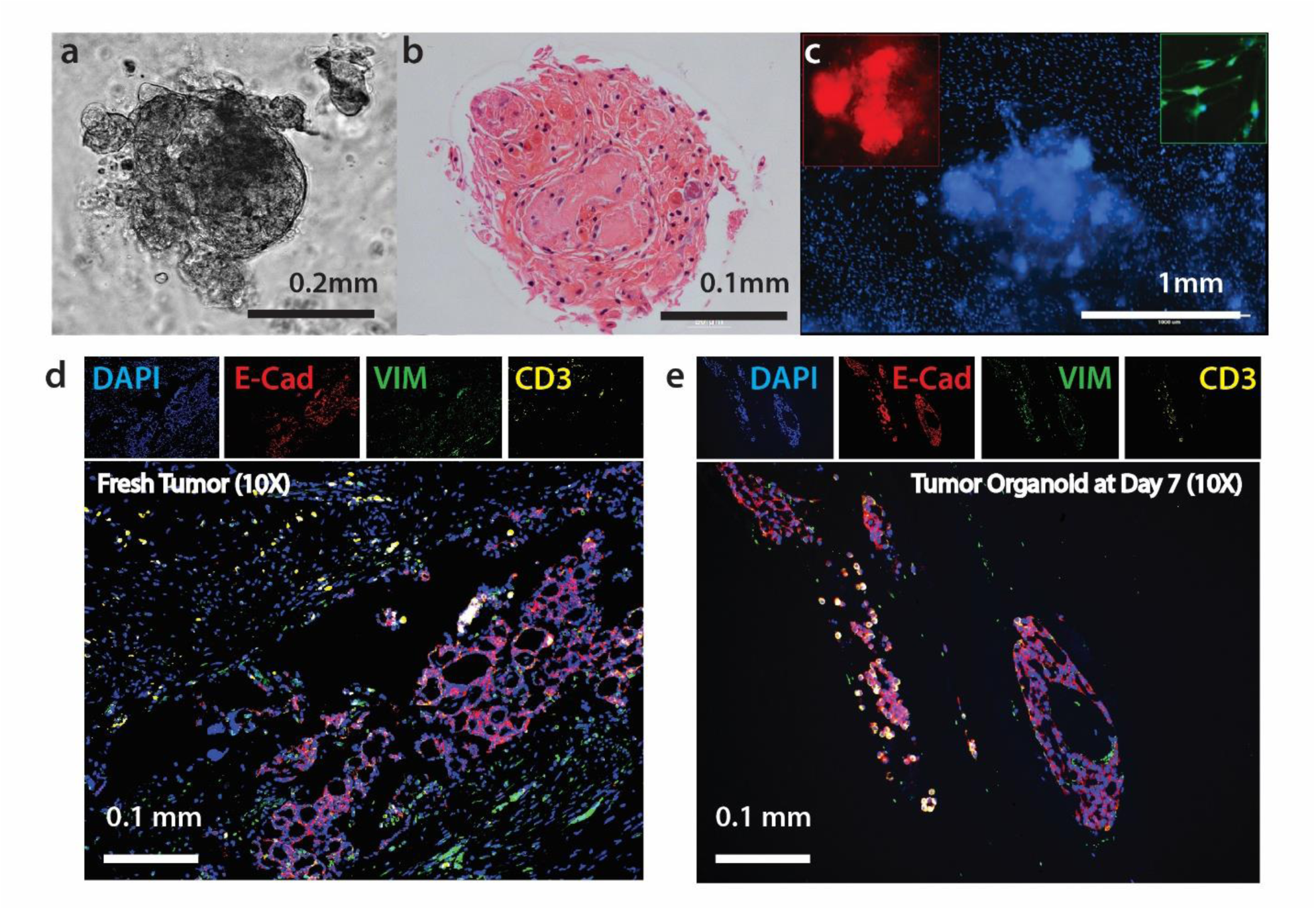
Characterization of head and neck tumor organoids grown from tumor tissue. (**a**) Brightfield microscopy and (**b**) histological section show squamous cell carcinoma tumor organoids maintain heterogeneous tissue structure. (**c**) Tumor organoid labeled with nucleus staining dye (Hoechst 33342). Inset: magnified view of cancerous tissue (red: DiI lipophilic tracer) and fibroblast rich stroma (green: DiO lipophilic tracer) around the organoid culture from two different organoids. (**d, e**) Fluorescence immunohistochemistry comparing (**d**) fresh tumor sample and (**e**) tumor organoid grown from adenoid cystic carcinoma sample labeled with four markers: blue showing DAPI (nuclei); red, E-Cadherin (tumor epithelial cells); green, Vimentin (tumor-associated fibroblasts); and yellow, CD3 (tumor-infiltrating T-cells).

### RLM enables imaging of glucose metabolism in tumor organoids

In contrast to normal cells, cancer cells largely avoid the mitochondrial tricarboxylic acid cycle and instead rely on aerobic glycolysis for energy production. The advantage of aerobic glycolysis, also known as ‘the Warburg effect’, is still not clear but the phenomenon is the molecular basis for assessing tumor burden *in vivo* using FDG-PET. Several studies using optical metabolic imaging have shown that tumor organoids display a glycolytic profile similar to that of solid tumors^10,25,26^, however, the FDG uptake of tumor organoids has not yet been examined.

Using a single-photon sensitive microscope, we imaged FDG uptake in organoids derived from clinical tissue specimens (Supplementary Fig. 2). Organoid cultures were incubated with FDG (1 mCi/ml) at 37°C for 1-2 h to give the radiotracer enough time to diffuse into the 3D tissues and be metabolized, then washed for 30 min. The organoids were then gently dissociated from their matrix by slow micro-pipetting, mounted onto a 0.5 mm-thick CdWO_4_ scintillator, and imaged using RLM as previously described.^23^

A few adjustments were made to the standard RLM protocol to allow thick 3D organoid samples to be imaged. RLM images are constructed from the radioluminescence flashes which originate from individual positron decays occurring inside the organoid tumor. In one approach, called digital imaging, individual scintillation flashes collected by the EMCCD are precisely counted and converted to the number of radioactive decays to yield a composite map^27^ (Supplementary Fig. 3). Although this digital approach is the method of choice for 2D cell imaging, it is less efficient for 3D tissue samples due to the presence of a large number of cells within a small volume. While problematic for digital imaging, the high count-rate creates enough scintillation signal for fast and direct analog measurements of the whole organoid using a single camera exposure of 10-300 s, no pixel binning, and an electron-multiplication gain of 600/1200. We further use a 20X objective (NA =0.75) to achieve an imaging field of view of 1.5 mm and an image pixel size of 3.2 µm.

For accurate quantification, we obtained a calibration curve from a known radioactivity distribution (Supplementary Fig. 4). The obtained calibration data was found to be concordant with the exponential decay curve of ^18^F, down to very low activity and camera signal. To verify the accuracy of this calibration method, we compared the radioactivity of organoids estimated from the images to the total radioactivity of individual organoids measured using a gamma counter.

To quantify radiotracer concentration (Bq/pixel) in the oPEM images, the calculated total radioactivities were scaled according to the signal intensities of the individual pixels. An intensity colormap of the radioluminescence intensity depicts the distribution of FDG inside the tumor organoids and reveals spatial metabolic heterogeneity (Fig 3a & Supplementary Fig. 5). The combination of brightfield and radioluminescence images plays a similar role as clinical PET/CT images, where FDG accumulation can be assigned to a specific region or tissue structure with high glucose metabolism.

**Figure 3.**
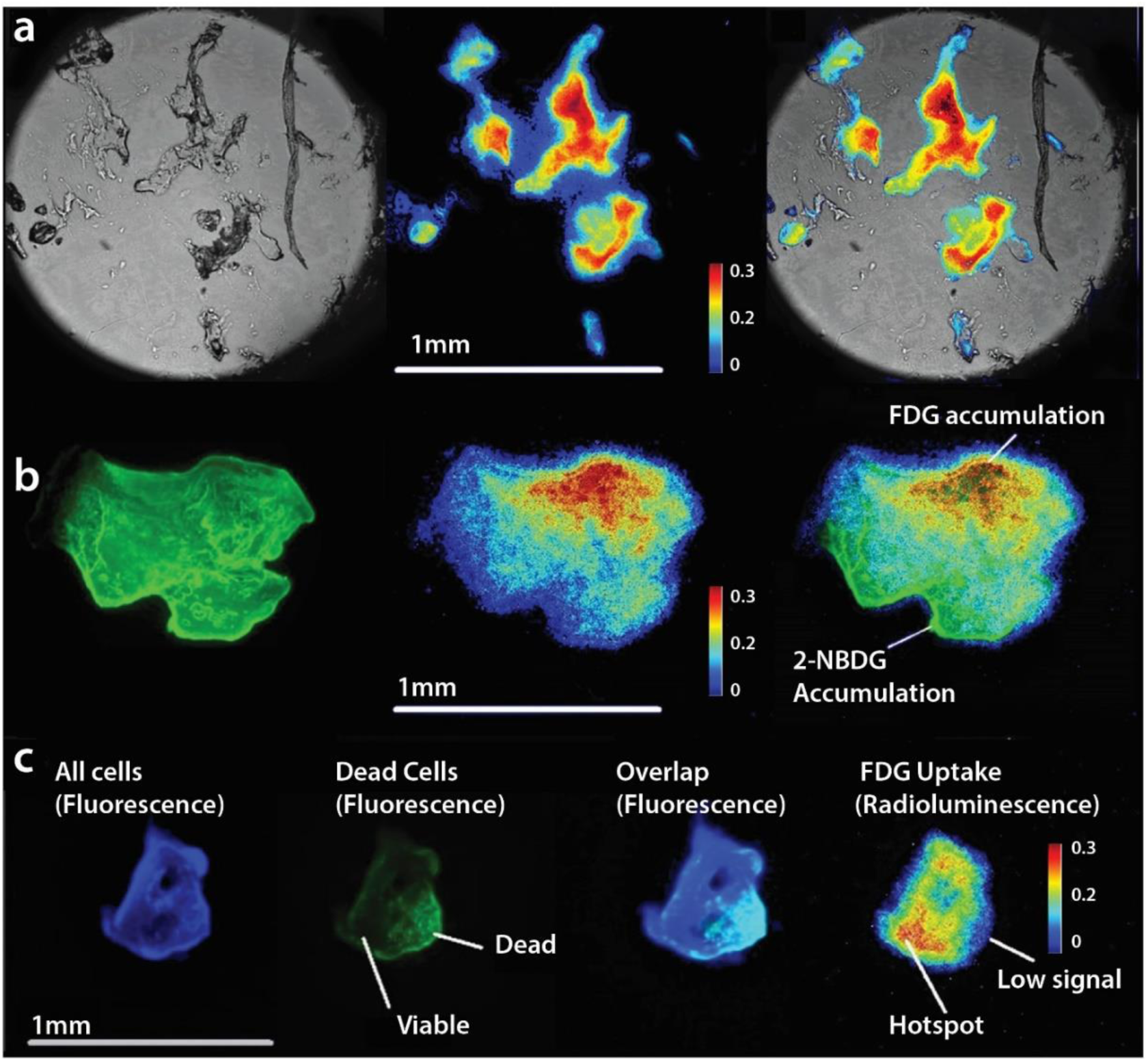
Positron emission microscopy of tumor organoids. (**a**) Brightfield image (*left*), radioluminescence image (*middle*) and overlay (*right*) show the correlation between FDG uptake distribution and tissue structure. FDG uptake is elevated in a majority of the organoid. (**b**) A comparison between fluorescence imaging with 2-NBDG, a fluorescent glucose analog (*left*) with oPEM imaging with FDG (*middle*) indicates inconsistent co-localization (*right*) of two probes. (c) Tumor organoid labeled with live/dead fluorescent stains (3 left panels) shows that FDG uptake (right panel) is associated with tissue viability.

A comparison with 2-(N-(7-Nitrobenz-2-oxa-1,3-diazol-4-yl)Amino)-2-deoxyglucose (2-NBDG), a fluorescent analog of glucose, revealed that FDG was more specifically taken up by tumor organoids, indicating the superiority of FDG over 2-NBDG to probe cellular metabolism (Fig. 3b). It is worth mentioning that, unlike FDG, any fluorescent analog of glucose would require a bulky fluorophore group attached to the carbon backbone. This requirement significantly alters the biochemical properties of glucose, making it less reliable to study glucose metabolism in cells and tissues. Besides, the fluorescent signal is sensitive to several extrinsic parameters such as polarity, molecular crowding, pH, and interaction with the environment resulting in unwanted fluorescence enhancement and quenching.

From this perspective, developing microscopic imaging techniques using FDG would be extremely useful for accurate imaging of glucose metabolism. Unlike other metabolic imaging approaches, FDG uptake has a clear biological meaning, as its uptake represents the total flux of glucose into glycolysis. Additionally, compared to fluorescence imaging, information obtained from FDG-based imaging is relevant to clinical practice and can be used to correctly predict or compare *in vivo* glucose metabolism in tumors using FDG-PET. The approach can also be extended to other PET tracers for which there is no equivalent fluorescent analog available.

In another set of experiments, tumor organoids were co-labeled with live/dead fluorescent probes to assess the specific localization of FDG uptake (Fig. 3c). Generally, the radioluminescent hot spots were localized within the viable regions of the organoids. This positive correlation indicates that FDG uptake is specific and representative of metabolically active regions. No uptake was observed in non-viable organoid tissue, indicating that FDG is not retained unless metabolized by live cells. However, it should be mentioned that fluorescent live/dead probes are not expected to correlate exactly with FDG uptake, as their uptake is based on the difference of membrane permeability between live and dead cells instead of their metabolic state.

Finally, the spatial resolution of oPEM is crucial to elucidate the spatial distribution of PET tracers (e.g. metabolic heterogeneity) inside complex tissue structures. Due to the high specificity of the tracer uptake, the high resolution of oPEM could be advantageous to differentiate cancerous tissue from stromal counterpart while judging therapeutic responses in these organoids. The full-width half-maximum of a tiny structure, measured using oPEM, was found to be 17 ± 1 µm (Supplementary Fig. 6). This result suggests that the resolving capacity in the current experimental condition is very high when the sample is placed very close to the scintillator surface. The resolution may decrease in thicker organoids, but it is still many folds higher than the > 5 mm resolution of clinical PET imaging. To achieve even higher spatial information, organoids could be cut into thin sections and imaging could be performed digitally (as described before), although this invasive approach would prevent kinetic and longitudinal imaging.

### Quantification of glucose metabolism inside organoid tumors

The emergence of patient-derived tumor organoids calls for new tools to produce reliable, reproducible, and quantifiable biomarkers of therapeutic responses that are consistent with clinical practice. To highlight the clinical relevance of organoid imaging, we quantified FDG uptake into organoids and, using clinical FDG-PET data, the tumors from which they were derived. A key step in this analysis is to identify an intrinsic parameter that represents the metabolic state of the tissue, independent of the spatial dimensions of the system, the amount of FDG available to tumor cells, and the time course of the uptake. The standardized uptake value (SUV) is a well-established metric that accounts for the size of the patient and the injected tracer dose, but not for the time kinetics of the tracer in the plasma compartment. The SUV may therefore not be a suitable metric to compare organoids and in vivo tumor uptake, since there is no tracer clearance for organoid cultures.

This study considers instead the net uptake rate *K*_*i*_, a metric derived from Patlak analysis assuming irreversible tracer uptake. The coefficient *K*_*i*_ has units of inverse time and represents the net transport and trapping of FDG into tumor cells as a fraction of the known concentration of FDG in the plasma compartment. Since virtually all clinical PET scans are performed as static scans, a simplified analysis was conducted. First, we assumed the y-intercept (*V*_D_) to be negligible compared to the tumor uptake. The approximate *K*_i_ rate computed using this approach, sometimes referred to as the fractional uptake rate (FUR), is valid for scans acquired at late time points. Second, in place of arterial blood sampling, we used a standardized input function derived from a population of over 101 patients.^28^ The model provides an input function adjusted for patient height and weight. Third, we assumed that the volume of blood in the region of interest was negligible compared to the volume of tumor tissue. Using this model, we computed the *K*_i_ of tumors from FDG-PET images, and that of organoids from oPEM and/or gamma counting measurements.

Generally, FDG concentration in the organoids was on the order of 10,000 kBq/ml, much higher than the uptake measured in tumors using PET (∼10 kBq/ml). This apparent discrepancy arises due to the difference in the concentration of FDG available to tumor cells. Although the FDG dose used for organoid imaging (∼30 MBq) was lower than the injected doses in patients (∼300 MBq), the final concentration of the tracer was much higher in the organoids culture medium (∼30, 000 kBq/ml) than in the patient plasma (∼30 kBq/ml) due to the vastly different distribution volume for the two biological systems. It is interesting to notice that despite these differences, the calculated *K*_*i*_ values fall in the same order of magnitude for both *in vivo* tumors and *in vitro* organoids (Supplementary Table 2).

To test whether FDG-oPEM captures inter-patient variability, we cultured organoids from two patients who each presented with thyroid nodules (containing papillary thyroid carcinoma) of contrasting FDG avidity. The first patient (T1) presented with an FDG-avid thyroid nodule (SUV=8.7; Supplementary Fig. 7) whereas the second patient (T2) had a FDG-cold nodule, with no visible uptake in the PET scan (SUV=1.8; Supplementary Fig. 8). The difference in metabolic intensity between the two nodules was reproduced *in vitro* in organoids derived from the same tumors using oPEM. Organoids from the first patient showed significantly higher FDG influx (*K*_*i*_=1.0 ± 0.3 %/min) than those from the second patient (*K*_*i*_=0.7 ± 0.2 %/min).

However, it could be argued that the difference in FDG activity may be influenced by differences in the genetic background and disease stage of the different patients. To overcome this confounding factor, we established organoids from a unique patient (T3) who presented with two thyroid nodules of similar size with contrasting FDG findings (Fig. 4a). The first nodule, located in the left lobe of the thyroid, had intense FDG uptake (SUV=32.4) whereas the second nodule, located in the right lobe, was cold and not visible on the PET scan (SUV=2.5; Supplementary Fig. 9). Both nodules were pathologically confirmed to be papillary thyroid carcinoma after resection. Thus, we had two different tissue samples of the same disease in the same patient to test whether metabolic heterogeneity is conserved when the tumor cells are grown as organoids. We calculated the FDG influx rate (*K*_*i*_) into the nodules and found it to be 3.0 %/min for the left side and *K*_*i*_=0.23 %/min for the right side. Similarly, oPEM imaging revealed a stark difference in FDG uptake between two groups of organoids (Fig. 4b). Although there was metabolic heterogeneity among the organoids derived from the same source, organoids derived from the patient’s left nodule had 10-fold higher FDG influx rate (*K*_*i*_ = 1.4 ± 0.3 %/min) on average than those derived from the right nodule (*K*_*i*_ =0.04 ± 0.02 %/min; Fig. 4c). Overall, the data from 4 tumors suggest that the bulk metabolic state of solid tumors is conserved in organoids *in vitro* (Fig. 4d); in other words, oPEM can functionally recapitulate the original PET imaging of the patient performed in the clinic.

**Figure 4.**
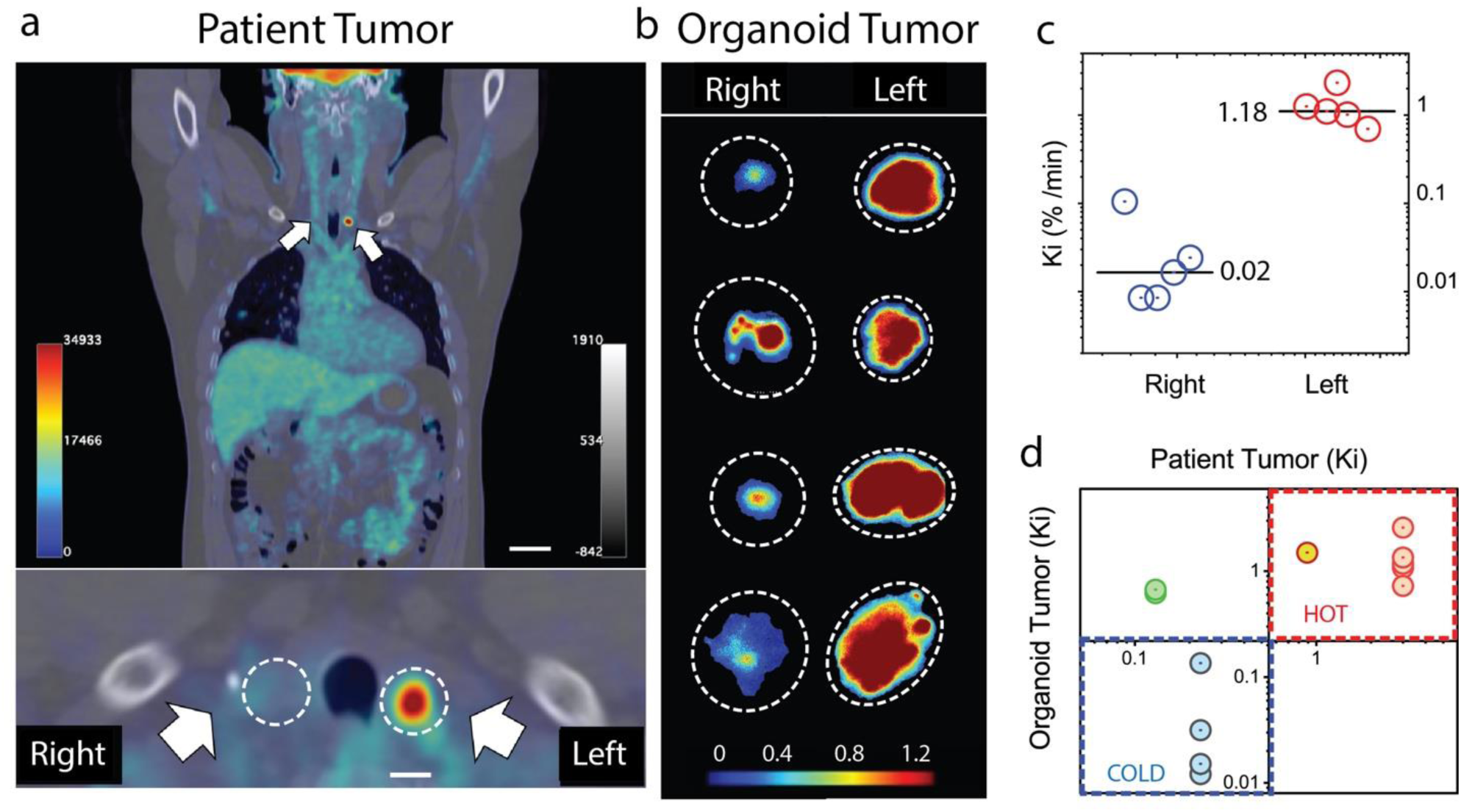
Comparison of PET vs oPEM imaging (**a**) PET/CT scan of patient T3. Two papillary thyroid carcinoma nodules are indicated by white arrows and show contrasting levels of FDG uptake. (**b**) oPEM images of tumor organoids derived from these nodules. The organoids retained the contrasting metabolic identity of the original nodule. The dotted lines show the extent of the individual organoids. (**c**) Scatter plot of FDG influx constant (*K*_*i*_) for organoids derived from left and right nodules. The median is labeled and shown by a black solid line. The organoids from the left nodule have more than 10 times higher mean uptake than those from the right nodule. (**d**) Scatter plot showing the correlation (Pearson’s *r*= 0.756, *P* < 0.005) between the *K*_*i*_ of organoids (n=13) and *K*_*i*_ of the tumor of origin (n=4).

### Assessing drug response in tumor organoids

Given this association between *in vitro* and *in vivo* metabolism, we sought to investigate the ability of oPEM to replicate, in an *in vitro* setting, the use of FDG-PET for monitoring therapeutic response in patients. The metabolic response of head-and-neck organoids was assessed using FDG-oPEM after cisplatin treatment (0, 1, and 10 µM for 24 hours; Fig. 5a & Supplementary Fig. 10). For clinical relevance, dosages of 1 & 10 µM were chosen (the IC50 value and local concentration of cisplatin in patient tumors often fall near that range^29^). FDG uptake shows a dose-dependent decrease in glucose metabolism after treatment with cisplatin. In quantitative terms, the uptake of FDG was 315±24 Bq, 118±16 Bq, and 12±10 Bq for the three dosages of cisplatin, respectively (Fig. 5b). These values correspond to approximately ∼3 MBq/ml, ∼1 MBq/ml, and ∼0.1 MBq/ml, respectively. The significant decrease in metabolic response within 24 h of drug treatment reflects the high sensitivity of this approach. Unlike cell viability assays, this method can identify cell fate early on the basis of cell glycolysis. This is the very same reason why FDG-PET has proved to be more effective than anatomical measurements for monitoring therapeutic response in patients. Fig. 5a also shows the live/dead fluorescent co-labeling, which was performed to identify the viable regions of the organoids. Again, most of the FDG uptake was within the viable regions, indicating an inverse relationship between dead cell staining and FDG signal through the organoid structure (Fig. 5c). However, a perfect correlation between viability and FDG uptake was not expected as the dead cell stain does not label the low-metabolic viable cells.

**Figure 5.**
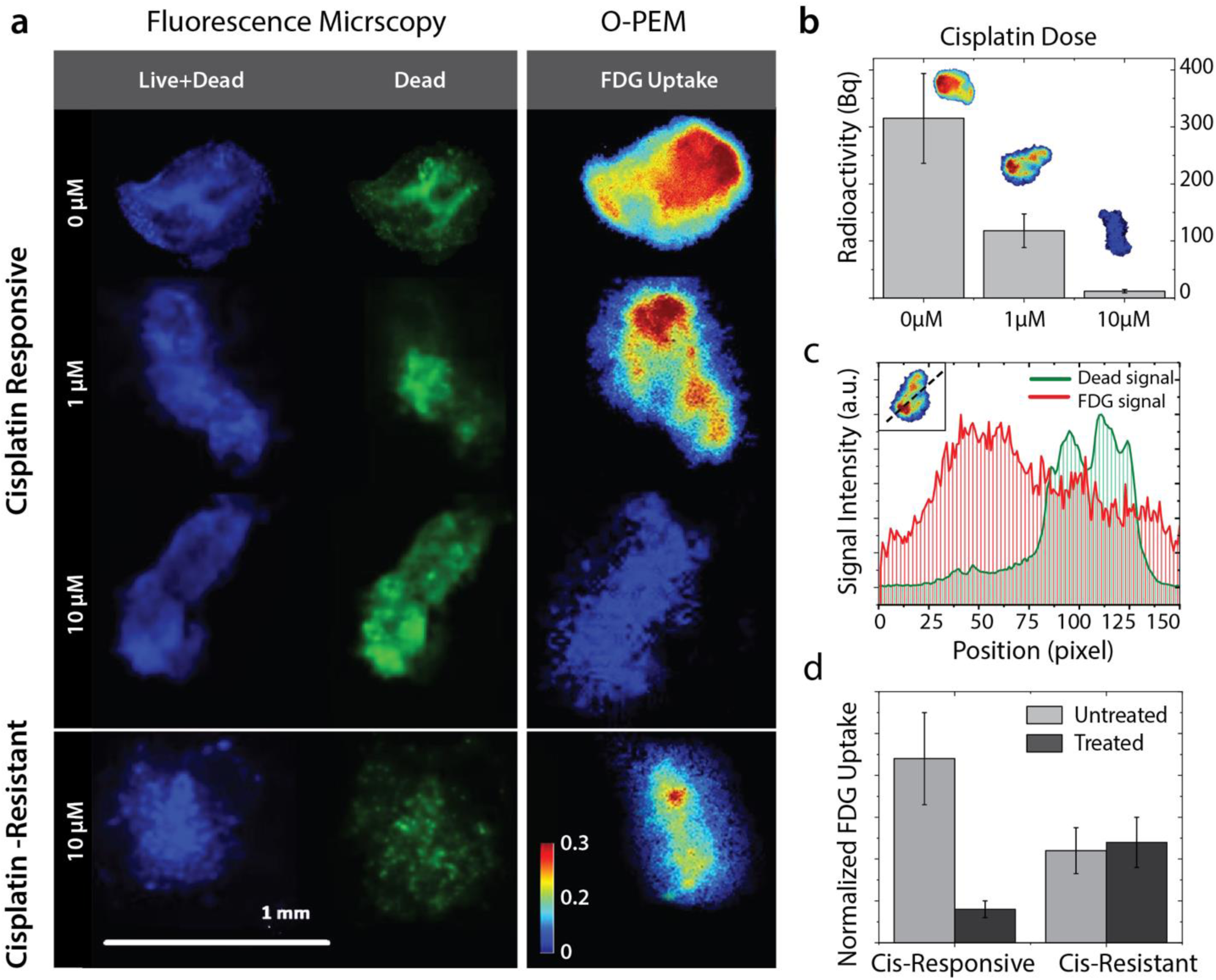
oPEM & fluorescence co-analysis of metabolic activity after cisplatin treatment. (a) Untreated organoid shows a spatial pattern of cellular viability (fluorescence stain; left panel) consistent with the FDG uptake profile (right panel). Generally, organoids treated with increasing doses of cisplatin experience a decrease in their metabolic activity. However, organoids obtained from a cisplatin-resistant patient maintained substantial metabolic activity despite cisplatin treatment. (b) Quantitative FDG uptake inside tumor organoids treated with 0, 1, and 10 µM dose of cisplatin. (c) The intensity profile of the FDG signal and green channel along the dotted line (inset) shows a spatial anti-correlation. (d) Comparison showing normalized FDG uptake for cisplatin-responsive and cisplatin-resistant organoids.

Finally, to investigate the potential of oPEM for predicting clinical responses, we grew tumor organoids from a patient (S3-CR) who had relapsed after cisplatin and radiation treatment. Given the poor response of this tumor to cisplatin, we hypothesized that organoid models would retain this resistant phenotype *in vitro*. Organoids from this patient grew slowly and had low FDG uptake, even at baseline, compared to other organoids. Cisplatin treatment (10 µM) did not substantially change the metabolism of the organoid at the 24 h timepoint (lower panel, Fig. 5a & Supplementary Fig. 11), suggesting that resistance to cisplatin was maintained *in vitro*. A normalized uptake index was calculated as the ratio of the FDG uptake to the amount of labeled DNA in the organoid (Fig. 5d). Taken together, our data indicate that oPEM imaging could be useful to predict clinical responses by screening tumor organoids obtained from individual patient tumors and identify drug resistance ahead of treatment.

## Discussion

In summary, a major finding of our study is that clinical PET tracers, such as FDG, can be imaged with high spatial resolution after they distribute in 3D organoid models of cancer. Using oPEM, we find that tumor organoids recapitulate specific metabolic features of the tumors of origin. The method can also monitor drug response of organoid cultures. This concept is analogous to the use of FDG-PET for monitoring treatment response in cancer patients. The method is safe, cost-effective, relatively fast, and capable of screening samples with 100-fold higher resolution than a clinical PET scan. Thus, oPEM could be a useful tool for preclinical studies based on organoid models.

oPEM is an extension of RLM specifically adapted for imaging 3D tumor organoids. While originally developed for imaging 2D cell monolayers, we found that RLM can readily image thicker specimens with only a moderate loss of spatial resolution. The importance of 3D cancer models is well recognized for modeling the unique features of the microenvironment of solid tumors, making these models relevant for several applications. Organoids can be grown from freshly excised tumor tissue and assembled into panels for testing novel therapeutics. They can also be used to predict the response of individual patients to therapy. Yet, despite their growing popularity, organoids have not yet been imaged with clinical PET imaging agents. Results from this study show that RLM can perform the equivalent of a PET scan on patient-derived organoids to reveal the heterogeneous metabolic state of the tissue. Non-viable tissue does not retain FDG, indicating that FDG uptake is a specific surrogate for the flux of glucose into cells. In addition, our analysis of a small dataset of matched PET/organoid scans suggests that the metabolic state of tumors is preserved when these tumor cells are grown *in vitro* as organoids. Accordingly, FDG imaging of organoids could serve an endpoint to assess the response of individual organoids to therapy. One advantage of oPEM is that it can help bridge the gap between *in vitro* tumor models and clinical trials in patients by providing a unified biomarker (FDG uptake) for rapid translation of knowledge from the pre-clinical to the clinical arena.

The use of organoid models is expanding rapidly, driven by factors such as cost, throughput, robustness, and versatility. Many of the hallmarks of cancer—metabolic reprogramming, drug resistance, and even metastasis—can be modeled using organoid models^26,30^. In this context, the use of oPEM would be beneficial for studies that are challenging to perform *in vivo*. For instance, oPEM could help elucidate the onset of metabolic reprogramming during tumorigenesis in normal tissue organoids^31,32^, which could have value for understanding the feasibility of detecting early-stage tumors with PET. Moreover, oPEM could be used in areas beyond oncology to image normal tissue organoids and study-specific disease pathologies^4,32,33^. PET tracers are being developed for a wide range of diseases, ranging from neurological disorders to cardiovascular and infectious diseases. Organoids will provide a versatile platform to accelerate the development of improved PET tracers for these emerging applications.

In the clinic, PET does not allow clinicians to predict which treatments are most likely to elicit a response; rather, the information is obtained post-hoc, weeks after the treatment has started, thus delaying the switch to more effective therapy. There is an urgent need for approaches that can identify suitable therapies, especially for patients who fail first-line therapy. A number of recent studies support the use of tumor organoid models to predict the response of individual patients to therapy. Vlachogiannis et al. generated a live organoid biobank from patients with metastatic gastrointestinal cancer who had previously been enrolled in phase I or II clinical trials and found that the organoids had similar molecular profiles to those of the patient tumor^30^. Yao et al. have shown that tumor organoids can predict chemoradiation responses of locally advanced rectal cancer, reinforcing their value as a companion diagnostic tool in rectal cancer treatment^34^. An *in vitro* test, developed by Ooft et al. based on tumor organoids from metastatic lesions, accurately predicted response to irinotecan monotherapy or 5-FU–irinotecan combination therapy, but failed to predict 5-FU–oxaliplatin combination therapy outcome^6^. According to a study by Scognamiglio et al, patient-derived organoids can predict individual responses to immunotherapy in patients with low or no immunohistochemical PD-L1 expression^35^. Thus, in this context, oPEM could become a valuable tool to provides a way to investigate the same clinical biomarker of PET to predict therapeutic response in a noninvasive fashion. This capability could be also used as part of co-clinical trials, which use organoid models derived from patients enrolled in clinical trials.^30^

PET is an outstanding tool for imaging molecular processes in vivo. Its picomolar-concentration sensitivity is unequaled in the clinical arena and countless radiotracers are available to image the unique hallmarks of cancer^36,37^. A new generation of PET tracers is now entering clinical use that can image cancer-specific targets such as bombesin receptors, prostate-specific membrane antigen (PSMA), somatostatin receptor, α_v_β_6_ integrin, and newly identified fibroblast activation protein inhibitors (FAPI)^38-40^. Some of these emerging tracers are unique in that they do not target tumor cells but, rather, specific alterations of the tumor microenvironment. For instance, tumor-specific markers are expressed on the neovasculature of tumors^41^. Others, such as FAPI, are displayed by tumor-associated fibroblasts. Additionally, tracers have been developed to assess the activity of tumor-infiltrating lymphocytes^42^. However, traditional 2D cultures lack a suitable microenvironment to evaluate these tracers *in vitro* under realistic conditions. Tumor organoids contain a broad spectrum of cells including immune and stromal cells, and thus can be used as a model to evaluate and screen advanced PET tracers. We, therefore, expect tumor organoids to become a valuable model to test and improve oncologic PET tracers.

It should be noted that RLM can image different types of charged particles over a wide range of energies, and thus it is not limited to any specific radioisotope such as ^18^F. Commonly used PET radioisotopes include ^18^F, ^68^Ga, ^64^Cu, ^124^I, and ^11^C, all of which can be easily imaged with high resolution. The different positron range of these isotopes only minimally affects spatial resolution because only a small section of the scintillator is in focus by the microscope. Additionally, therapeutic alpha and beta radiation could be imaged for microdosimetric evaluation of radiopharmaceutical therapy and response assessment in organoid tumor models. Organoids may, therefore, provide a compelling new platform for the assessment of theranostic regimens.

There are several advancements of oPEM that could further enhance organoid imaging. First, as the current study is limited by the small number of patients, a larger-scale study would be necessary to advance this method towards its intended application. Secondly, tomographic imaging capabilities would be useful to localize the tracer in 3D, in thick specimens. These capabilities have long been available for optical imaging thanks to the development of confocal microscopy and other approaches. However, tomography is more challenging to achieve for RLM, as, in the current technique, the light is collected not from the object but from the scintillator plate which absorbs the ionizing radiation. In theory, limited-angle tomography could be achieved if the 3D angle of incidence of individual positrons could be measured. A third potential improvement would be to develop a high-throughput technology to measure FDG uptake in hundreds of organoids for large-scale drug screening and other applications. Various prospective drug screening platforms have been developed using 2D cell culture, spheroid culture, mouse & zebrafish avatars, and patient-specific induced pluripotent stem cells ^13,17,43-45^. Similar to these, a high-throughput organoid screening would allow testing a range of treatment options for individual patients, but with the right balance between accuracy, ease of use, turn-around time, and cost-effectiveness.

In conclusion, we presented the use of oPEM for imaging tumor organoids. Given the expanding use of organoid models in research, we envision a number of emerging applications for this imaging technique. For instance, FDG-oPEM could be used to select suitable therapies for individual patients on the basis of FDG imaging of patient-matched organoids. Second, the approach could be used as an endpoint to screen drug candidates using heterogeneous panels of organoids. Finally, oPEM could be used as part of co-clinical trials, which are conducted using samples collected during clinical trials. The development of small-animal PET scanners 15 years ago has demonstrated the utility of developing preclinical imaging tools that have strong relevance to the clinical workflow. Accordingly, oPEM could become a standard technology for imaging tumor and normal tissue organoids using PET tracers, thereby expanding the use of organoids models for translational research.

## Methods

### Human specimen collection

Freshly resected primary and metastatic tumor tissues were obtained through the Stanford Tissue Bank from patients undergoing surgical resection at Stanford University Medical Center. All experiments utilizing human material were approved by the Institutional Review Board. Written informed consent for research was obtained from donors before tissue acquisition.

### Organoid culture

Excess fat or muscle tissue from human tumor tissue was removed to enrich the samples for epithelial cells. Tumor tissues were minced finely (approximately 2-3 mm) on ice, washed twice in DMEM/F-12 (Invitrogen) containing 1X Primocin (InvivoGen) and followed with ACK Lysing Buffer (Thermo Fisher Scientific) to remove red blood cell contamination. Minced tissues were washed again with DMEM/F-12 media and resuspended with cold matrigel (Cultrex® reduced growth factor basement membrane matrix, type 2, Trevigen). Droplets of approximately 40 μl were plated on the bottom of pre-heated 24 well culture plates (E&K Scientific). After plating, culture plates were incubated at 37°C for 30 minutes to let the BME solidify. Organoid culture media (EN media: EGF+Noggin) containing DMEM/F-12 supplemented with 10% Noggin-conditioned media, Nicotinamide (10 mM, Sigma), N-Acetylcysteine (1 mM, Sigma), B-27 without vitamin A (1X, Invitrogen), Pen-Strep (1X, Invitrogen), and EGF (50 ng/mL, R&D Systems) was added. The medium was changed every 2-3 days and organoids were passaged every 2-4 weeks by dissociation with 1 ml of TrypLE Express (Life Technologies) at 37°C for 5 min. Tissue suspension was sheared using 1 ml pipette. Digestion was monitored closely to prevent excess incubation in trypsin. Organoid pellets obtained after centrifugation at 1500 rpm were washed with DMEM/F-12 media (containing 10% FBS and Pen-Strep) and replated at a 1:3 split with BME matrigel. For Cisplatin (Millipore Sigma) treatment, 1-10 µM of Cisplatin was added to the media for 24 h with DMSO as a control. For cryopreservation of organoid tissues, freezing media (90% FBS containing 10% DMSO) was used with a standard freezing protocol. The organoids shown in figure 2e was cultured using a slightly modified technique to better retain the tumor-infiltrating immune cells as described in a previous study by Neel J et al.^5^

### Histology and immunofluorescence analysis

Organoids were fixed in 4% paraformaldehyde, paraffin embedded and sectioned by Stanford Human Pathology and Histology Service Center. Sections (4-5 μm) were deparaffinized and stained with hematoxylin and eosin (H&E). For immunofluorescence staining, the following primary antibodies: anti-E-cadherin (BD, 610181, 1:1000), anti-vimentin (Millipore, AB1620, 1:1000), and anti-CD3 (Dako, A0452, 1:100). All secondary antibodies were used at 1:500. All images were captured on a Zeiss Axio-Imager Z1 with ApoTome attachment.

### Radiolabeling and sample preparation

For imaging of glucose metabolism using combined fluorescence and radioluminescence microscopy (RLM), organoids were incubated in glucose-free DMEM medium for 1 h before addition of FDG (1 mCi/ml). FDG was produced at the Stanford radiochemistry facility using an on-site cyclotron. The organoids were incubated with FDG for 1-2 h followed by washing with phosphate buffer saline (PBS) thrice with gentle rocking. The hydrogel was gently disrupted and the organoids were picked using a pipette tip and transferred to PBS for final washing. The organoids were washed thrice in PBS to remove all unbound FDG. Individual organoids were carefully collected and gently mounted on a CdWO_4_ scintillator plate (0.5 mm thick), which itself was mounted onto a standard glass coverslip (0.1 mm thick, Fisher Scientific). The specimens were then promptly placed on the microscope stage for multimodal imaging. The specimen remained hydrated during the short imaging time.

### Instrumentation and imaging

Several scintillator materials can be used for RLM. CdWO_4_ has a moderately high light yield (12,000–15,000 photon/MeV), high effective atomic number (Zeff = 64), high density (7.9 g/cm3), and no significant afterglow. The set-up was mounted on a custom-built wide-field microscope equipped with short focal tube lens (4X/0.2 NA; Nikon, CFI Plan Apochromat λ), 20X/0.75 NA air objective (Nikon, CFI Plan Apochromat λ), and deep-cooled electron-multiplying charge-coupled device (EM-CCD; Hamamatsu Photonics, ImagEM C9100-13).

Brightfield images were acquired with no EM gain. For RLM imaging, images were taken with a 20X objective, an exposure time of 10-300s, an EM gain of 600/1200, and 1×1 pixel binning. We used the brightfield mode to set the microscope into focus. Optimal radioluminescence focus was achieved when the organoids displayed sharp positive contrast in the corresponding brightfield image. For fluorescence microscopy, we used 387 mm/447 mm filter set (Semrock, filter ref: DAPI-1160B-000) for Hoechst imaging (live marker) and 469 nm/525 nm filter set (Thorlabs, filter ref: MDF-GFP1) for 2-NBDG or SYTOX green (dead marker) imaging.

### Quantification of FDG from oPEM

To quantify the FDG dose inside organoids, the EMCCD radioluminescence signal was calibrated with a known amount of radioactivity. Radioluminescence signal from a microdroplet of FDG with known activity was measured with a 5 min exposure time. Images were acquired over 8 h while the radioactivity gradually decayed down to a very low value. The intensity of the signal in a region of interest was plotted against time and compared with the decay curve of FDG. The camera signal integrated over the microdroplet area was scaled linearly to obtain the spatial concentration of radioactivity in Bq/pixel.

### Influx rate computation

To properly account for the time-varying concentration of FDG available to tumor cells, we calculated the rate of FDG influx into patient and organoid tumors (*K*_*i*_). The parameter *K*_*i*_ was calculated from a simplified Patlak model (assuming zero-intercept) with the following equation:

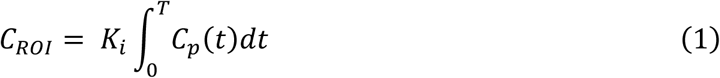

where *C*_*ROI*_ is the concentration of the tracer in the region of interest and *C*_*p*_ is the concentration of tracer in the plasma. *C*_*p*_ was assumed to be constant in the organoid tumor as they were incubated in a fixed concentration of FDG for a known duration. For patients, the estimation of *C*_*p*_ is more complex and ideally would necessitate serial arterial sampling to determine a time-dependent input function. To overcome this challenge, we used a standardized input function (SIF) derived from a population of 101 patients^28^ to compute a patient-specific input function *C*_*p*_:

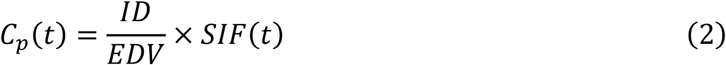

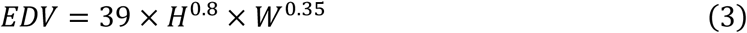

where *ID* is the injected FDG dose, *EDV* is the estimated distribution volume of FDG in a patient of height *H* and weight *W* estimated using equation (3), and *SIF*(*t*) is the function provided by Shiozaki et al.

The FDG concentration *C*_*ROI*_ was estimated from the maximum SUV value in the patient tumors. In organoids, *C*_*ROI*_was calculated by measuring the total radioactivity in a gamma counter (Hidex) and dividing by individual organoid volume. The volumes were estimated from the area measured using optical microscopy, assuming a spherical shape of those organoids. A calibration grid (Thorlabs, cat. no. R1L3S3P) was used for accurate estimation of the organoid area (cross-section).

## Acknowledgments

This work was funded in part by NIH grants R01CA186275, U19AI116484, U01DK085527, U01CA217851, U01CA176299, U01DE025188, and K00CA223019. The authors thank the Stanford radiochemistry facility for providing FDG. They also wish to gratefully acknowledge assistance from Tae Jin Kim for assistance using the radioluminescence microscope.

## Author Contributions

SK performed multimodal PEM imaging, developed new methodologies, analysed data and wrote the initial manuscript. JHS performed organoid culture and histology of all patient-derived tumor samples. JBS and JEN provided surgical tumor samples. In addition, JBS supervised organoid development. NC performed a comparative analysis between fresh tumor and organoid culture under CK’s supervision. VF processed and analysed clinical PET/CT data for *k*_i_ calculation. GP conceived the research idea, developed *k*_i_ analysis method, wrote the manuscript and supervised the study.

## Data availability

The main data supporting the results in this study are available within the paper and its Supplementary Information. The raw EMCCD and PET–CT data are available from the corresponding author on request.

## Notes

### Competing Interest Statement

The authors have declared no competing interest.

