## Supplementary Figures and Tables for "High-resolution positron emission microscopy of patient-derived tumor organoids"

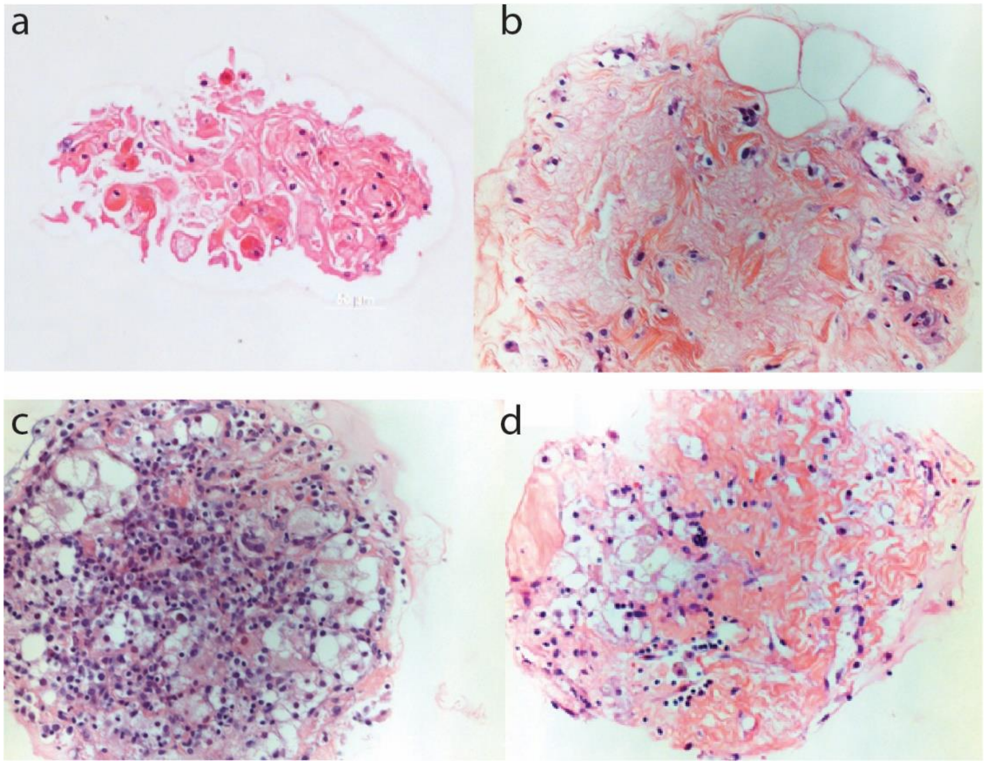

**SI Figure 1.** Histological imaging of patient-derived tumor organoid of (a) squamous cell carcinoma from Patient S1 and (b-d) papillary thyroid carcinoma from patient T1 shows heterogeneous tissue structure (20X magnification).

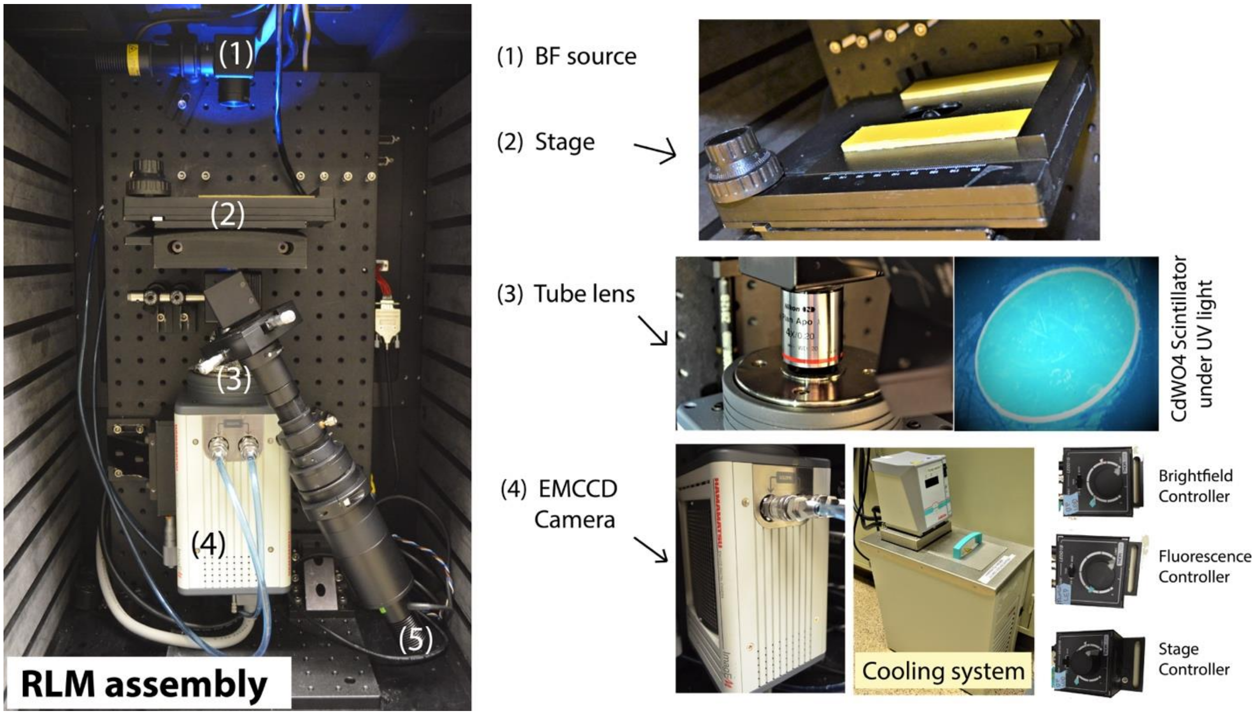

**Figure S2.** Overview of the microscope. Left, a photograph of the fully assembled microscope, showing (1) brightfield illumination, (2) stage assembly, (3) tube lens connected to a (4) EMCCD camera with a cooling unit, (5) LED light source for epifluorescence imaging. The optical train is composed of two microscope objective lenses aligned back to back. A 50-mm-focal-length lens (Nikon CFI Plan Apochromat λ 4×) is used in place of the standard f = 200-mm tube lens to demagnify the image and increase its brightness.

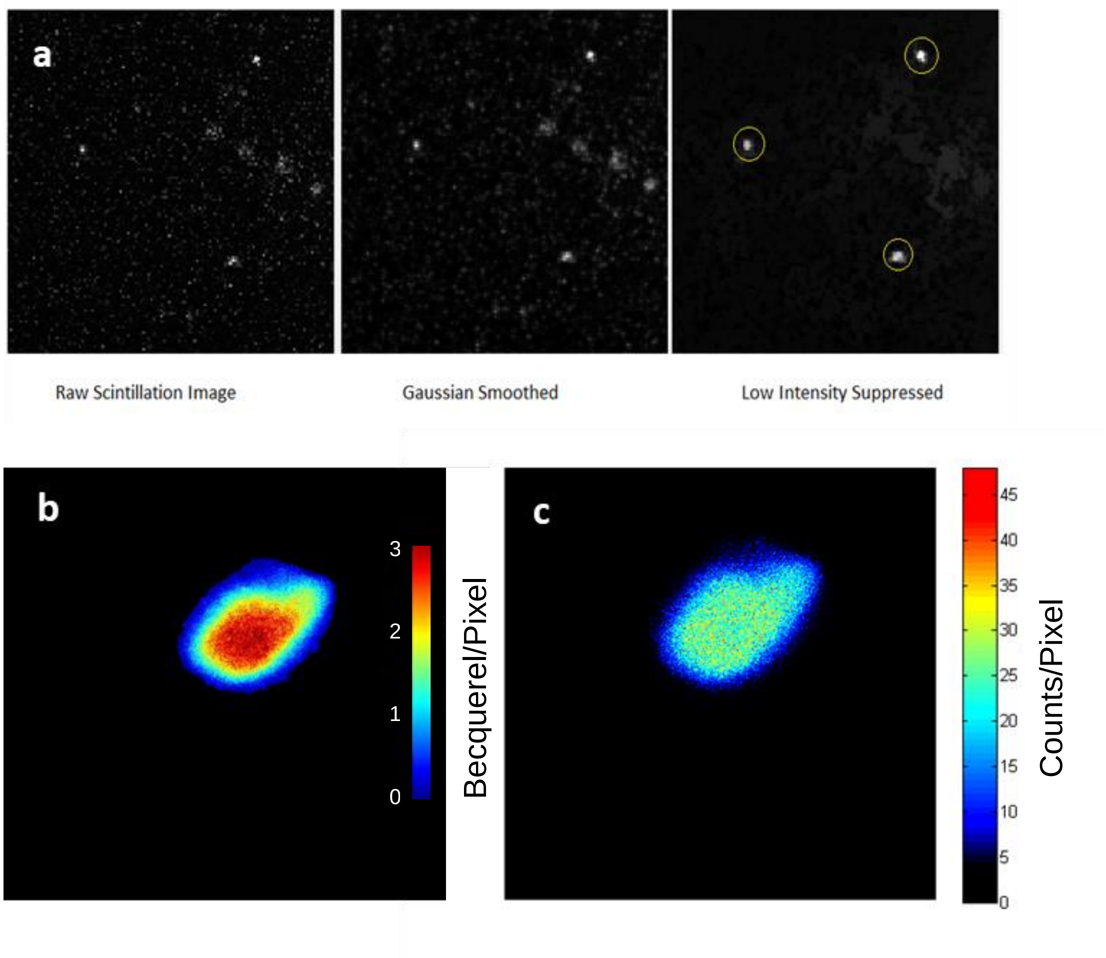

**Figure S3.** Optical reconstruction of a single frame using ORBIT software package. (a) The serial image processing of the raw image (left) allows elimination of the background signal and reconstruction of three most reliable events from the frame. (b) Analog and (c) digital oPEM image of tumor organoid grown from patient T1

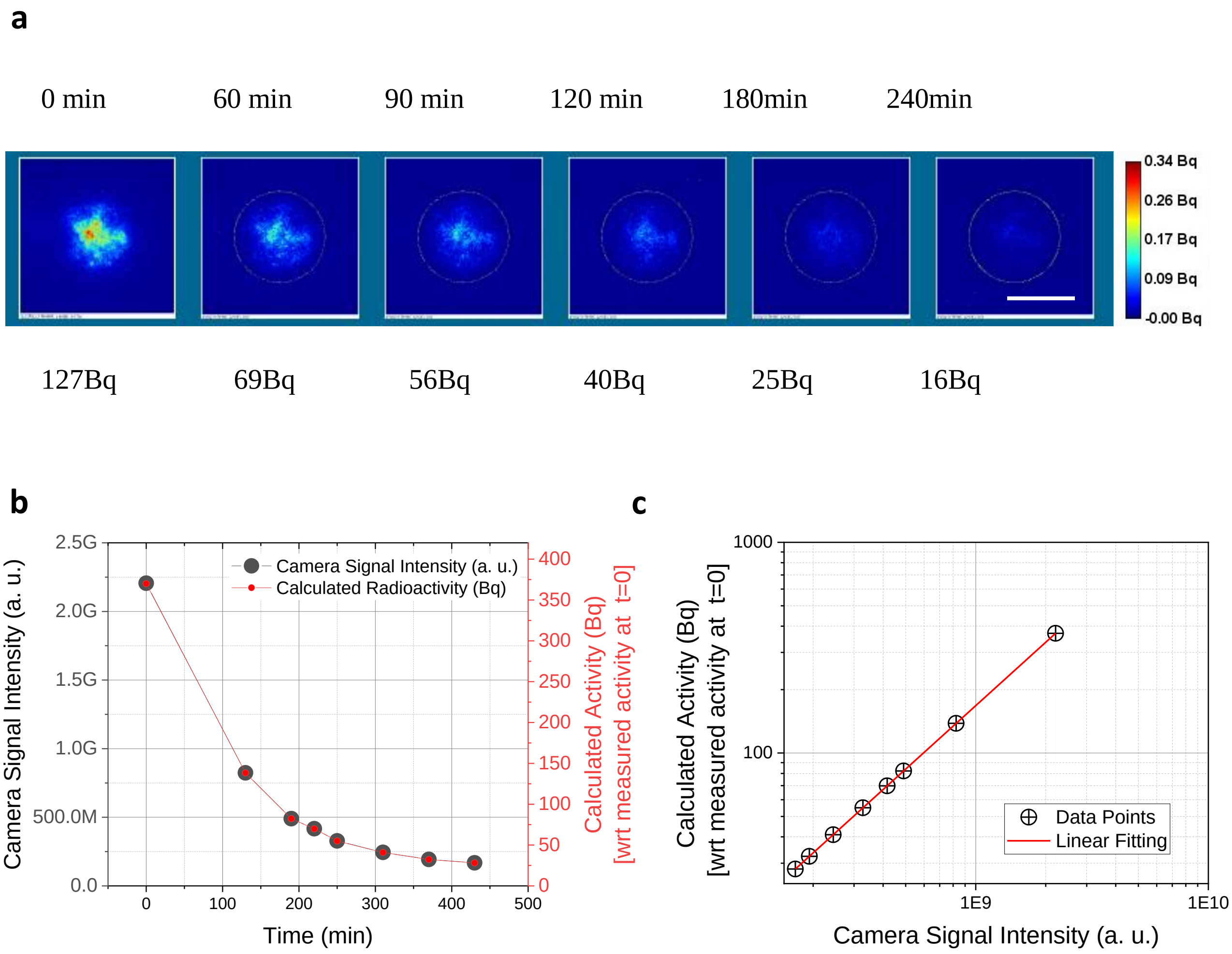

**Figure S4.** Quantification of radioactivity from oPEM image. (**a)** oPEM image of a small FDG droplet with known activity. Image taken at different time points shows decreasing radioactivity. Color bar: Bq/pixel (**b**) The time-dependent activity measured from the camera signal (within 1 $mm^2$) closely follows the decay-curve of $^{18}F$ (half-life ~110 min) (**c**) Calibration curve to convert camera signal into quantitative radioactivity measurements.

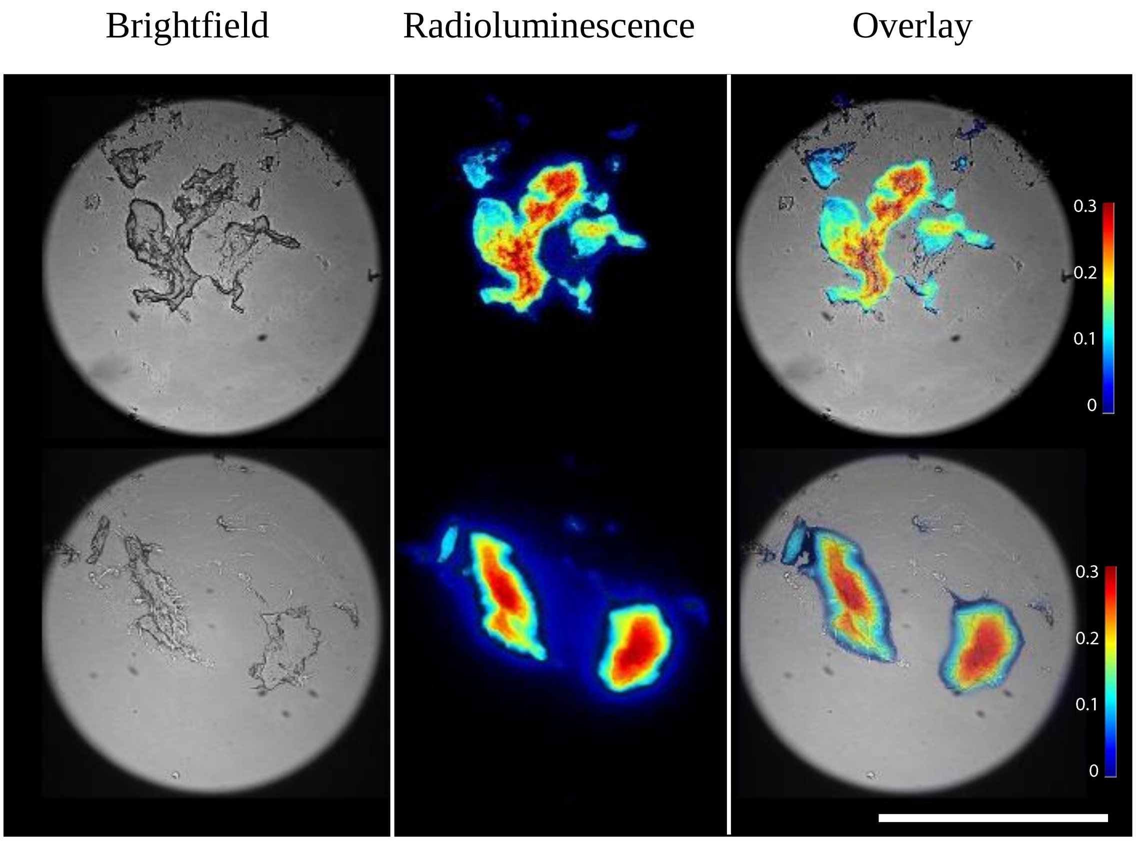

**Figure S5.** oPEM of tumor organoids derived from patient S1. Brightfield (left) and Radioluminescence (intensity colormap of FDG uptake, middle) shows certain hotspot indicating elevated metabolic activity in those part of the organoid structure in three different samples. An overlap (right) of bright filed image and color map reveals spatial distribution of FDG uptake in further detail. Scale bar is 100 µm. The intensity color bar (Color bar: Bq/pixel) shows increasing intensity from blue to red.

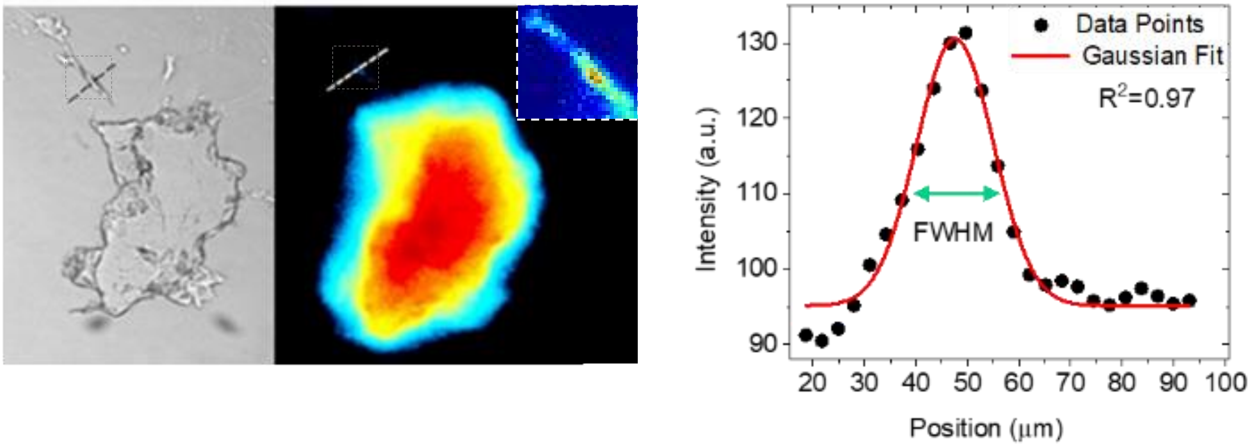

**Figure S6.** The spatial resolution of oPEM. Left: Brightfield and oPEM of an organoid (from Patient S1) showing a tiny feature. The inset shows a magnified image of the small structure. Right: Profile through radioluminescence image along the dashed line. FWHM was calculated to be 17.58 ± 0.84 μm along the dotted line

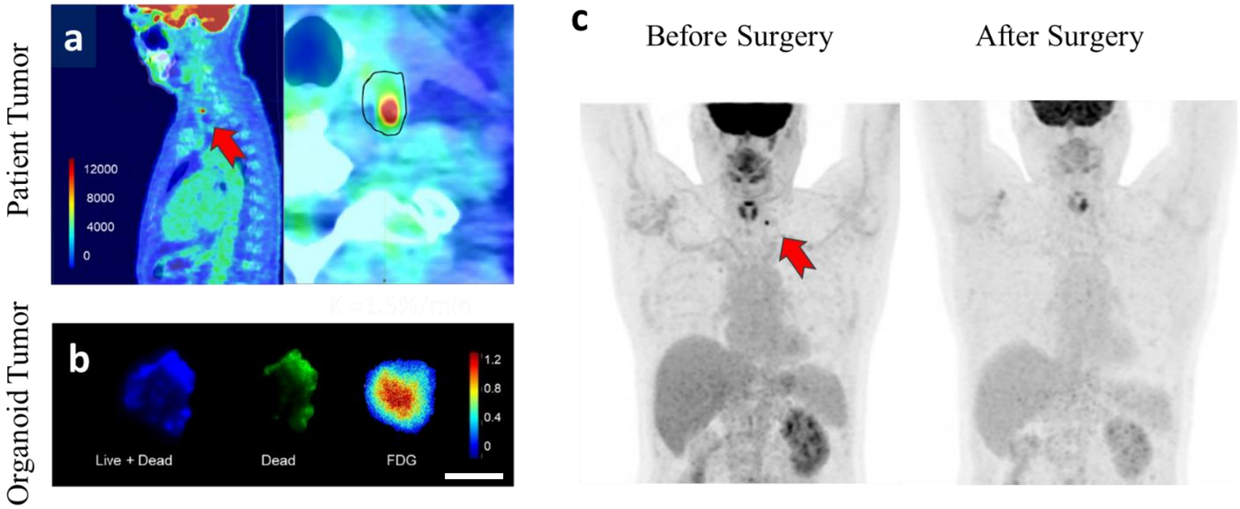

**Figure S7.** Comparison between FDG uptake in organoid and patient-of-origin tumors for patient T1. (A) PET image. A magnified view on the right panel shows a lymph node metastatic lesion from thyroid cancer. Scale bar: 10cm. Color bar: Bq/ml (B) oPEM imaging of tumor organoid grown from the same tumor of patient T1. Scale bar: 0.5mm. Color bar: Bq/pixel.

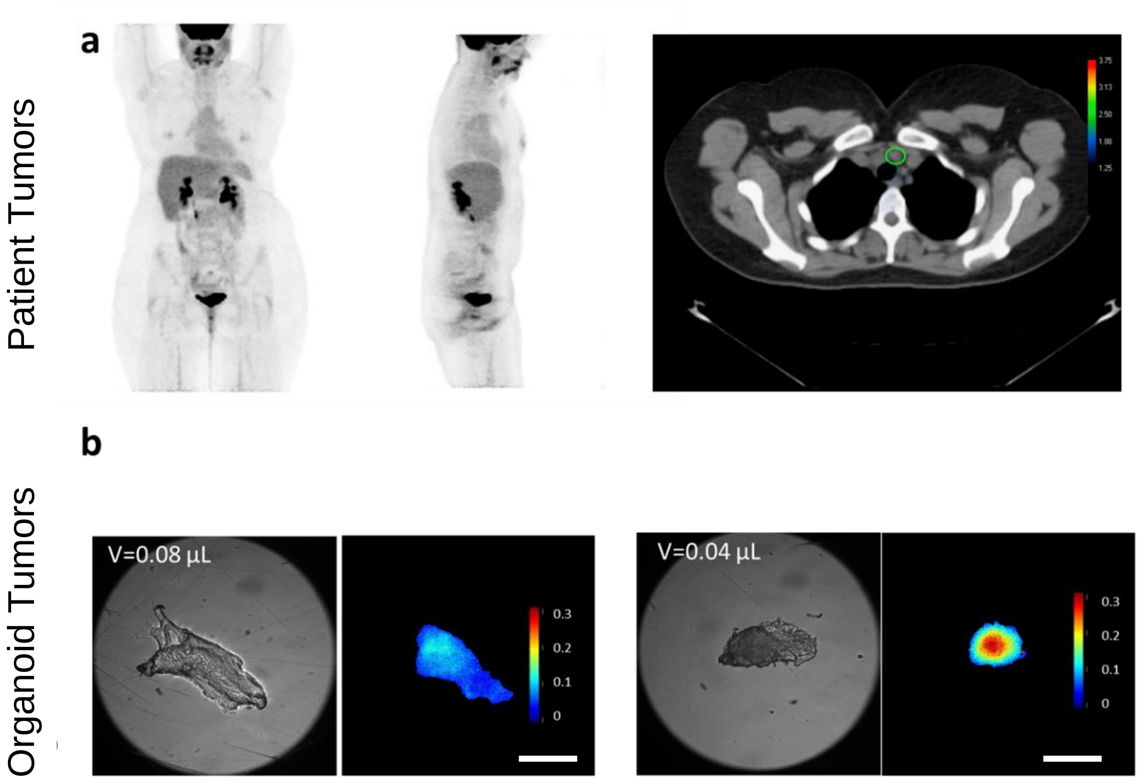

**Figure S8.** PET scan and oPEM of patient T2 (a) PET scan shows no visible uptake as the metastases were not FDG avid. (b) oPEM images show organoids derived from two of the excised lymph nodes metastases with moderate FDG uptake. Scale bar 1 mm.  Color bar: Bq/pixel.

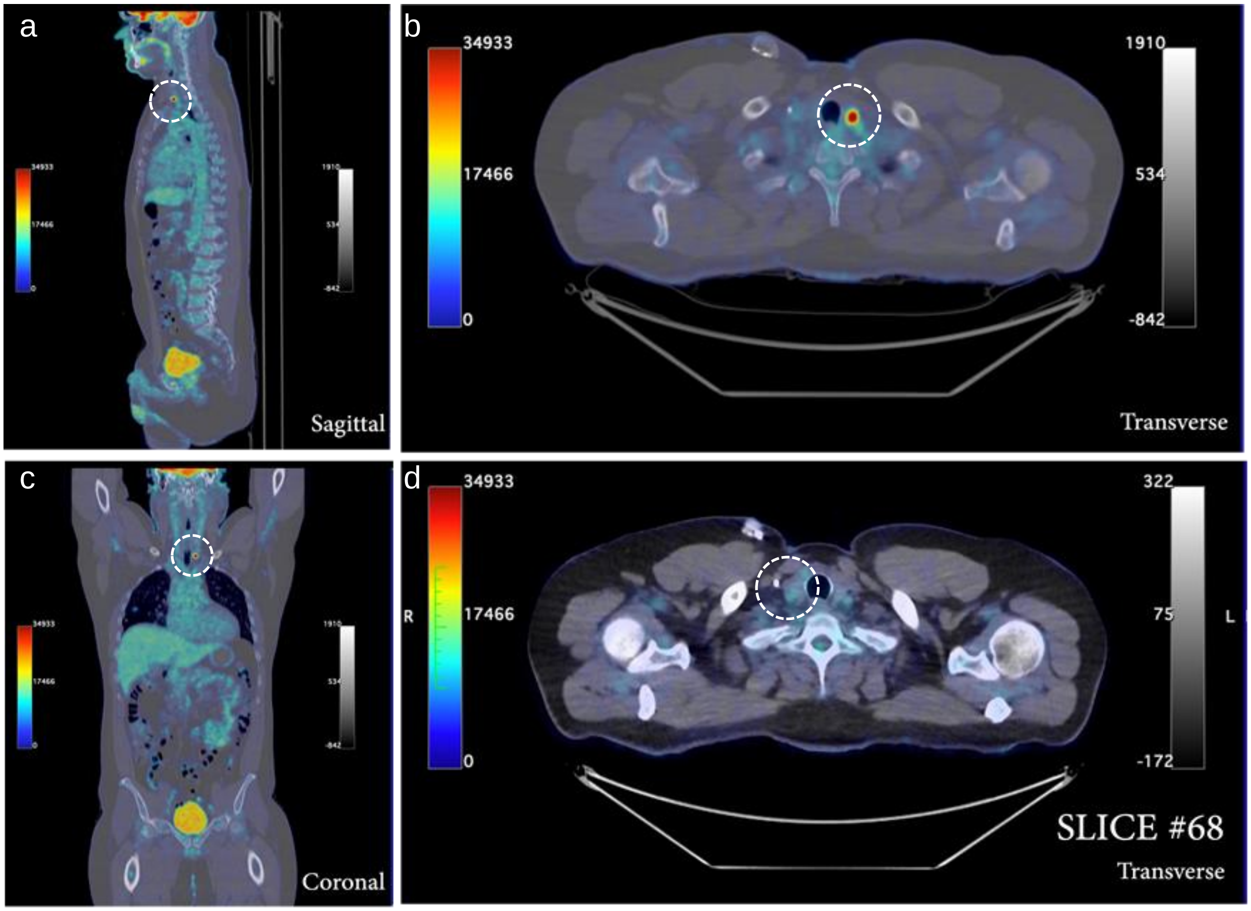

**Figure S9.** PET/CT images of patient T3. (a-c) A maximum intensity projection shows a hot nodule in the left side of the thyroid gland. (d) Slice #68 from the PET scan represents the location of the second thyroid nodule, of the right side of the thyroid gland, which did not take up any FDG.

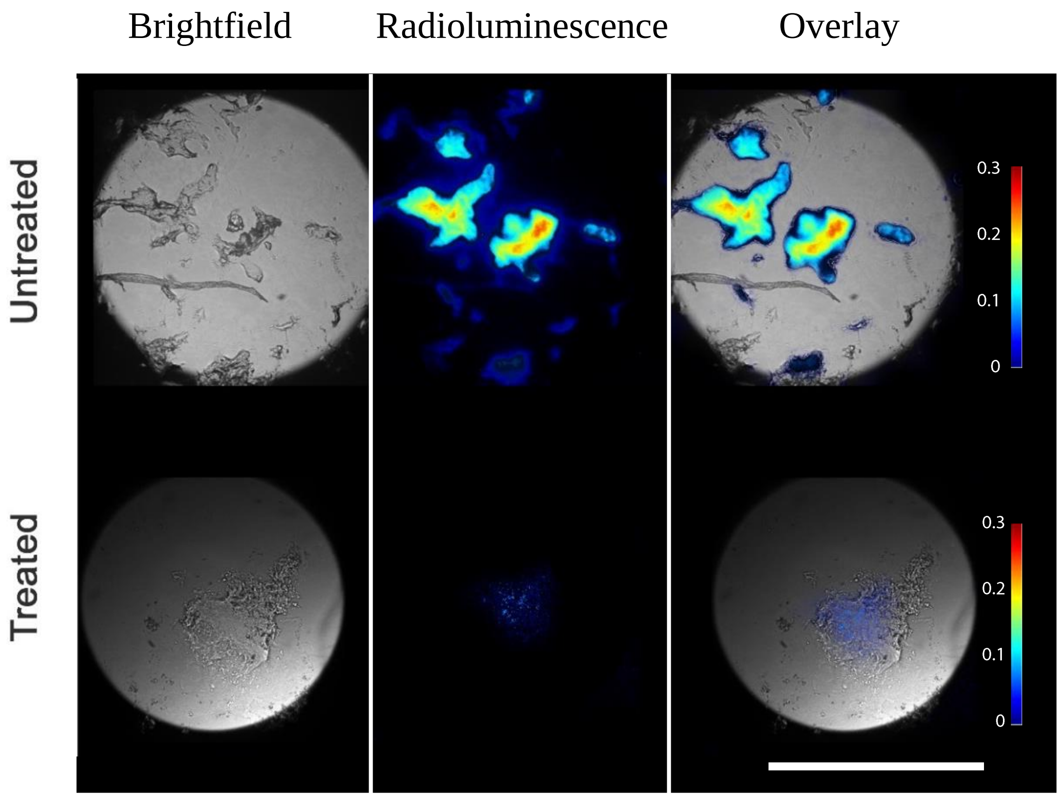

**Figure S10**. Comparison of oPEM images of organoids after 24h cisplatin treatment (10µM dose) and untreated organoids. The significant drop of FDG uptake indicates a decline in glucose in metabolism and cell proliferation. Scale bar: 1mm.  Color bar: Bq/pixel.

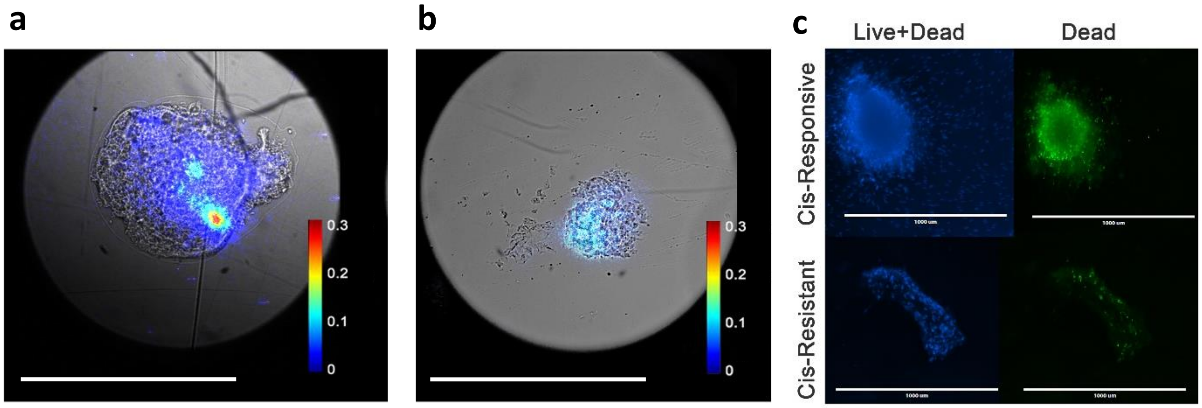

**Figure S11.** Imaging of PDOTs from a cisplatin-resistant patient (S3-CR). (a) Treated and (b) untreated cis-resistant PDOTs show equally low metabolic activity compared to the Cis-responsive PDOTs (c) Live/dead fluorescence labeling of cis-responsive and cis-resistant PDOTs after 10 µM cisplatin treatment. Scale bar: 1 mm. Color bar: Bq/pixel.

### Table S1: Patient and tumor details

| Patient ID | Gender | Age (y) | Height (cm) | Weight (kg) | Tumor type |
|---|---|---|---|---|---|
| **S1** | Female | 60 | 163 | 78 | Squamous cell carcinoma (pT4aN0, p16 negative). Tumor site: mandible, oral cavity |
| **S2** | Male | 70 | 165 | 59 | Squamous cell carcinoma (pT4aN3b, p16 unknown). Tumor site: tongue, oral cavity |
| **S3-CR** | Male | 48 | 177 | 75 | Squamous cell carcinoma (pT4aN0, p16 negative). Tumor site: base of tongue including lingual tonsil; oropharynx |
| **T1** | Male | 65 | 184 | 83 | Poorly differentiated papillary thyroid carcinoma (pT1a, N1b), metastasis to cervical lymph node |
| **T2** | Female | 29 | 172 | 106 | Papillary thyroid carcinoma (pT1b, N1b), diffuse sclerosing variant, with extensive lymphovascular invasion |
| **T3** | Male | 45 | 183 | 107 | Papillary thyroid carcinoma (cT1b, cN0, cM0). Brain metastasis. Sertoli cell tumor with lymphovascular invasion of right testis (rpT1a, pN2, cM0, S0) |

### Table S2: Pharmacodynamical parameters for the patient and organoid tumors

| | Patient ID | FDG Dose (MBq) | Distribution Volume (L) | Plasma Conc. (KBq/mL) | Target activity (KBq) | Target volume (mL) | Target Conc. (KBq/mL) | $K_i$ (%/min) |
|---|---|---|---|---|---|---|---|---|
| **Patient Tumor** | T1 | 340.4 | 297.7 | 1.1 | 9.9 | 0.4 | 19.8 | 0.89 |
| | T2 | 495.8 | 307.8 | 1.6 | 2.9 | 0.4 | 5.8 | 0.13 |
| | T3 (R) | 407 | 324.5 | 1.2 | 3.1 | 0.5 | 6.2 | 0.23 |
| | T3 (L) | 407 | 324.5 | 1.2 | 40.6 | 0.5 | 81.2 | 3.0 |
| **Tumor Organoid** | T1 | 37 | 0.001 | 37000 | 12.25<br>(0.8)[a]<br>(1.0)[a] | 0.00018<br>0.00006<br>0.00010 | 64,809<br>(12,986)<br>(23,400)<br>(51,183) | 1.46<br>(0.29)<br>(0.53)<br>(1.15) |
| | T2 | | | | 4.7<br>(2.6)[a]<br>2.9 | 0.00015<br>0.00007<br>0.00010 | 29,729<br>(39,500)<br>27,865 | 0.67<br>(0.88)<br>0.62 |
| | T3 (R) | | | | 0.3<br>0.1<br>0.2<br>3.1<br>0.3 | 0.00051<br>0.00046<br>0.00035<br>0.00052<br>0.00024 | 539<br>198<br>672<br>6,012<br>1,397 | 0.01<br>0.01<br>0.02<br>0.14<br>0.03 |
| | T3 (L) | | | | 2.6<br>5.4<br>2.6<br>22.5<br>9.8 | 0.00055<br>0.00010<br>0.00008<br>0.00020<br>0.00016 | 47,980<br>52,292<br>32,263<br>114,742<br>59,906 | 1.08<br>1.18<br>0.73<br>2.58<br>1.35 |

[a] Values in round brackets are estimated from oPEM images in the absence of gamma-counting data
